# Flagellar motility and the mucus environment influence aggregation mediated antibiotic tolerance of *Pseudomonas aeruginosa* in chronic lung infection

**DOI:** 10.1101/2024.10.25.620240

**Authors:** Matthew G. Higgs, Matthew A. Greenwald, Cristian Roca, Jade K. Macdonald, Ashelyn E. Sidders, Brian P. Conlon, Matthew C. Wolfgang

## Abstract

*Pseudomonas aeruginosa* frequently causes chronic lung infection in individuals with muco-obstructive airway diseases (MADs). Chronic *P. aeruginosa* infections are difficult to treat, primarily owing to antibiotic treatment failure, which is often observed in the absence of antimicrobial resistance. In MADs, *P. aeruginosa* forms biofilm-like aggregates within the luminal mucus. While the contribution of mucin hyperconcentration towards antibiotic tolerance has been described, the mechanism for mucin driven antibiotic tolerance and the influence of aggregates have not been fully elucidated. In this study, we investigated the contribution of flagellar motility towards aggregate formation as it relates to the diseased mucus environment. We found that loss of flagellar motility resulted in increased *P. aeruginosa* aggregation and tolerance to multiple classes of antibiotics. Further, we observed differential roles in antimicrobial tolerance of the *motAB* and *motCD* stators, which power the flagellum. Additionally, we found that control of *fliC* expression was important for aggregate formation and antibiotic tolerance as a strain constitutively expressing *fliC* was unable to form aggregates and was highly susceptible to treatment. Lastly, we demonstrate that neutrophil elastase, an abundant immune mediator and biomarker of chronic lung infection, promotes aggregation and antibiotic tolerance by impairing flagellar motility. Collectively, these results highlight the key role of flagellar motility in aggregate formation and antibiotic tolerance and deepens our understanding of how the MADs lung environment promotes antibiotic tolerance of *P. aeruginosa*.

**IMPORTANCE:** Antibiotic recalcitrance of chronic *Pseudomonas aeruginosa* infections in muco-obstructive airway diseases is a primary driver of mortality. Mechanisms that drive antibiotic tolerance are poorly understood. We investigated motility phenotypes related to *P. aeruginosa* adaptation and antibiotic tolerance in the diseased mucus environment. Loss of flagellar motility drives antibiotic tolerance by promoting aggregate formation. Regulation of flagellar motility appears to be a key step in aggregate formation as the inability to turn off flagellin expression resulted in poor aggregate formation and increased antibiotic susceptibility. These results deepen our understanding of the formation of antibiotic tolerant aggregates within the MADs airway and opens novel avenues and targets for treatment of chronic *P. aeruginosa* infections.

## INTRODUCTION

*Pseudomonas aeruginosa* is a Gram-negative bacterium frequently found in environments associated with human activity and is capable of causing a wide range of opportunistic infections (1). Individuals with muco- obstructive airway diseases (MADs) such as cystic fibrosis (CF), chronic obstructive pulmonary disease (COPD), and non-CF bronchiectasis (NCFB) frequently suffer from recurrent and chronic *P. aeruginosa* lung infections (2). Collectively, MADs represent the 3^rd^ leading cause of death worldwide, primarily owing to COPD and the increasing prevalence of bronchiectasis (3). MADs are characterized by the accumulation of dehydrated mucus within small airways. Stagnant mucus provides a unique, nutrient-rich environment for *P. aeruginosa* colonization that promotes the formation of bacterial community structures termed aggregates (4–6). These aggregates exhibit biofilm-like properties including enhanced tolerance and resistance to antibiotics and are associated with antibiotic treatment failure (7). While high dose inhaled antibiotic therapies show better efficacy and less toxicity than traditional delivery methods (e.g., oral and intravenous), chronic *P. aeruginosa* infections remain recalcitrant to antibiotic therapy and develop clinical resistance at high frequency (8). Antibiotic treatment failure remains a major cause of decreased quality of life and early mortality.

While genetically encoded antibiotic resistance remains a global health threat, we hypothesize that the failure of antibiotics to eradicate chronic *P. aeruginosa* lung infections is primarily due to antibiotic tolerance and persistence. Tolerance is a state where bacteria fail to grow in the presence of antibiotics, but still survive. Persistence occurs wherein a subpopulation of bacteria can withstand antibiotics for prolonged periods of time and often will not succumb to antibiotics (9). Understanding the mechanisms that contribute to antibiotic tolerance and persistence is paramount to the development of more effective treatment strategies.

In the transition from acute to chronic lung infection, clonal populations undergo significant diversification both genotypically and phenotypically. Isolates from chronic infection often exhibit a loss of acute virulence factors and increased antibiotic tolerance and persistence (10). While chronic infection often results in a remarkable array of diversity, there are some common traits that evolve in the MADs airway environment. One of the most frequent adaptations is loss of flagella-mediated motility (11–13). Flagellin is a potent pro-inflammatory TLR5 agonist and swimming motility has been shown to stimulate the formation of neutrophil extracellular traps, which are a potent tool of neutrophils to trap and destroy pathogens (14,15). It has been suggested that the loss of flagellar motility is an adaptation that may allow for host immune evasion (16,17). However, MADs are an inherently inflammatory disease, and the loss of flagella has not been shown to reduce immune activation or inflammation (12,18). Several studies have investigated the relationship between loss of flagella and antibiotic susceptibility of *P. aeruginosa* under laboratory conditions (19,20). For instance, a mutant lacking the flagellar hook protein (*flgE)* displayed altered biofilm structure and a reduction in gentamicin penetration of the biofilm (20). Another study showed that a *flgK* mutant, which lacks another component of the flagellar hook complex, was more tolerant to the clinically relevant antibiotic, tobramycin (19). Specifically, the authors showed that a *flgK* mutant more readily formed aggregates in an agar gel. Despite these observations the role flagella and flagellar motility on tolerance in the diseased mucus environment is poorly understood.

There are many factors that have been shown to contribute to antibiotic tolerance, ranging from the specific environment to bacterial encoded factors. In the context of MADs, we have previously shown that mucin and DNA content within the environment can shape tolerance to tobramycin (20). Within the MADs lung environment, *P. aeruginosa* has been shown to reside as multicellular aggregate biofilms (4), that are thought to be highly tolerant to antimicrobial therapies. However, the requirements for aggregation and events that lead to the formation of aggregates are poorly understood.

Here, we investigated the role of flagellar motility in antibiotic tolerance and aggregate formation in the context of a MADs-like mucus environment. We found that mutants of the flagellar machinery are significantly more tolerant to antibiotics and that aggregate formation directly correlates with antibiotic tolerance. Using single cell motility tracking, we uncovered differential roles of the flagellar stator complexes, MotAB and MotCD, in tolerance and aggregation. Our results also suggest that regulation of flagella is important for the genesis of aggregation, and subsequently antibiotic tolerance. Our results significantly increase our understanding of the requirements of aggregate formation and describe the contribution of flagellar motility to antibiotic tolerance in the context of the MADs airway environment.

## RESULTS

### Aggregation increases in a mucin concentration dependent manner and correlates with antibiotic tolerance

Our previous work has shown that antibiotic tolerance increases as a function of mucin concentration (21). It has been shown that charged polymers, including mucin, drives aggregation of *P. aeruginosa* (22,23); as such, we investigated the relationship between aggregation, mucin concentration, and antibiotic tolerance. To assess aggregation and antibiotic tolerance, we used the wild type (WT) laboratory strain mPAO1 grown in synthetic CF mucus media 2 (SCFM2), a medium that mimics the CF lung environment (24,25), and modulated the mucin content to assess how mucin concentration impacts tolerance and aggregation. We assessed tolerance by growing bacteria in SCFM2 with various mucin concentrations for 8 hours, followed by treatment with high dose tobramycin (300μg/mL) for 24 hours. Aggregation was assessed by confocal microscopy using bacteria that expressed the fluorophore, dsRed-Express2 (26). To capture aggregation within the luminal media rather than surface attached bacteria, all confocal images were taken at least 10μm above the bottom of the growth chamber surface. Aggregates were quantified using the Imaris image analysis software (Oxford Instruments) and the surfaces function; surfaces greater than 5μm^3^ were considered as aggregates (27).

Consistent with our previous work, we observed a mucin concentration dependent effect on tolerance, with survival to tobramycin increasing as mucin concentration increased (**Fig. 1A)**. Similarly, aggregate size increased significantly as mucin concentration increased (**Fig. 1B**, **top panels, 1C, and Fig. S1**). We also observed a mucin concentration dependent reduction in the proportion of the biomass that remained planktonic (**Fig. 1D**). These data demonstrate that aggregation positively correlates with antibiotic tolerance, and that increasing mucin concentration shifts the population to a more aggregated phenotype.

**Figure 1.**
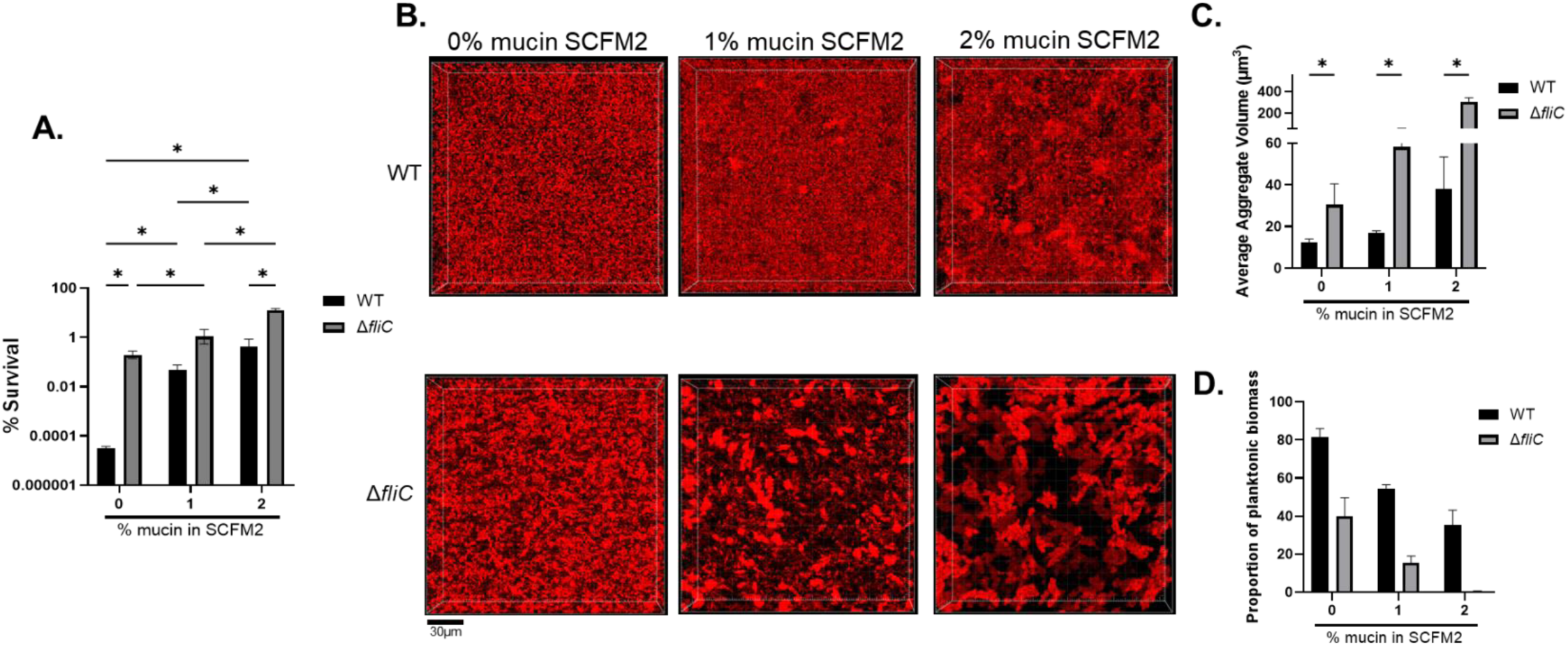
Aggregation strongly correlates with tobramycin tolerance in a mucin concentration dependent manner. **A)** WT mPAO1 or Δ*fliC* were grown in SCFM2 with the indicated mucin concentration for 8 hours, then treated with tobramycin (300μg/mL) for 24 hours. Percent survival is plotted as mean +/- SD. **B)** WT PAO1 or Δ*fliC* expressing fluorescent protein were grown in SCFM2 at various mucin concentrations for 6 hours prior to imaging via 3D confocal microscopy. Scale bar is 30μm. **C)** Quantification of aggregates from **B)** using Imaris. Aggregates were classified as surfaces >5μm3. Data represents the average +/- SEM aggregate volume from at least 3 independent images. **D)** Proportion of planktonic biomass (<5um^3^) from Imaris analyzed images. Planktonic biomass plotted as average +/- SEM from a minimum of 3 images. All data are representative of 3 independent experiments. \**P<0.05,* as determined by two-way ANOVA with Tukey multiple comparisons test **A**) or students *t-*test **C).**

### Loss of flagellar motility increases antibiotic tolerance to tobramycin and correlates with increased aggregation

One of the most common phenotypic adaptations observed in chronic *P. aeruginosa* isolates from MADs is the loss of flagellar motility (12,13). It was previously shown that flagellar mutants exhibit decreased susceptibility to tobramycin when embedded in an agar polymer gel (19). We investigated whether this trend existed in a more relevant mucus airway environment, and if similarly, aggregation of flagellar mutants were responsive to changes in mucin concentration. We utilized a non-polar deletion mutant of *fliC*, which encodes flagellin, the structural subunits of the flagellum fiber. A Δ*fliC* mutant was significantly more tolerant than WT mPAO1 to tobramycin and exhibited a significant increase in aggregation at all mucin concentrations tested. (**Fig. 1A-C**, **Fig S1**). Unsurprisingly, the proportion of planktonic biomass also decreased in Δ*fliC* compared to WT, with virtually no planktonic bacteria remaining in SCFM2 with 2% mucin (**Fig. 1D**). These data strongly suggest that aggregation may account for mucin concentration driven antibiotic tolerance. To confirm the role of the flagellum in tolerance, we also tested a non-polar *flgE* deletion mutant (Δ*flgE*). Both *flgE* and *fliC* mutants fail to produce flagellin or produce surface flagella (**Fig. S2A**). Similar to the *fliC* mutant, deletion of *flgE* resulted in a significant increase in tobramycin tolerance in SCFM2 with 2% mucin (**Fig. S2B**). Further, ectopic expression of *fliC* from its native promoter in the Δ*fliC* mutant (Δ*fliC*::ΦCTX-*fliC*) restored the tolerance to tobramycin to a level similar to WT (**Fig. S2C**). Based on the findings here, our previous findings, and its relevance to disease, we utilized SCFM2 with 2% mucin for the remainder of this study (21).

We next assessed whether the phenotypes of the *fliC* and *flgE* mutants were related to loss of the flagellum structure, or loss of motility. The flagellum is primarily powered by two stator complexes, MotAB, and MotCD. As such, we generated non-polar deletion mutants of the *motAB* and *motCD* stators. Deletion of either system alone or both stator complexes simultaneously does not prevent flagellin synthesis or the assembly of flagella on the cell surface (**Fig. S2A**). Like the non-flagellated mutants, deletion of *motCD* or both stator complexes (Δ*motABCD*) resulted in a significant increase in tolerance to tobramycin (**Fig. 2A**). In contrast, Δ*motAB* exhibited a decrease in tobramycin survival, suggesting a differential role of the stator complexes in antibiotic tolerance.

**Figure 2.**
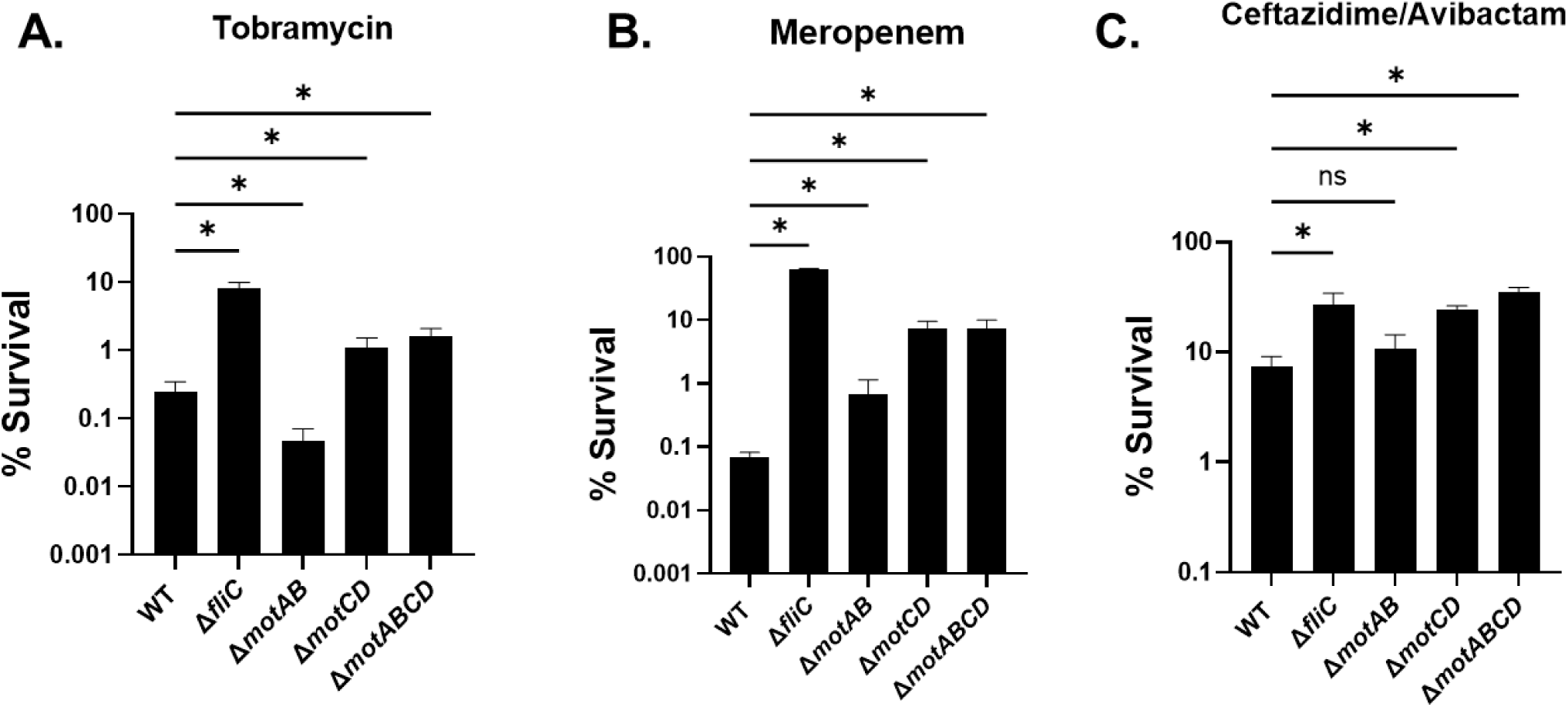
Loss of flagellar motility promotes tolerance to multiple classes of antibiotics. The indicated strains were grown in SCFM2 containing 2% mucin for 8 hours, then treated with **A)** 300 μg/mL tobramycin (aminoglycoside), **B)** 2000μg/mL meropenem, or **C)** 1000/40 μg/mL ceftazidime/avibactam for 24 h. *\*P<0.05* as determined by one-way ANOVA with Dunnett’s post-hoc test. Data shown are mean +/- SEM and are representative of 3 independent experiments.

While tobramycin is one of the most commonly used antibiotics for chronic *P. aeruginosa* lung infection, other antibiotics such as meropenem (carbapenems) and ceftazidime/avibactam (cephalosporin/β-lactamase inhibitors) are also used (28–30). Thus, we assessed whether loss of flagellar motility promoted tolerance to other classes of antibiotics. Similar to tobramycin, the Δ*motCD* and Δ*motABCD* mutants exhibited an increase in survival to both meropenem and ceftazidime/avibactam, similar to Δ*fliC* (**Fig. 2B, C**). Interestingly, the Δ*motAB* mutant showed either no difference in tolerance (ceftazidime) or a slight increase in survival (meropenem), further suggesting a differential role of the different stators in antibiotic tolerance.

The ability of bacteria to survive high concentrations of antibiotics for prolonged periods of time can promote resistance (31,32). We assessed the ability of flagellar mutants to withstand prolonged exposures to antibiotics by treating cultures for 72 hours (**Fig. S3**). We found that for all antibiotics tested, both WT and Δ*motAB* had similar susceptibilities while Δ*fliC,* Δ*motCD*, or Δ*motABCD* remained tolerant during prolonged antibiotic exposure.

### Flagellar stators differentially contribute to tolerance and aggregation

Given that non-flagellated mutants exhibit increased tolerance to several classes of antibiotics and showed increased aggregation, we investigated aggregate formation of the stator mutants. We found that the mutants that exhibited increased antibiotic tolerance (Δ*motCD* and Δ*motABCD*) also exhibited a significant increase in aggregate size compared to WT mPAO1 (**Fig. 3A-B)**. Consistent with the increase in aggregate size, there was a decrease in the proportion of planktonic biomass, indicating that more of the population was aggregated (**Fig. 3C**). The Δ*motAB* mutant, which was less tolerant to tobramycin, showed less aggregation than WT and exhibited a higher proportion of planktonic biomass. We also assessed traditional surface attached biofilm formation by the flagellar mutants in the presence of disease concentrations of mucin (2% w/v).

**Figure 3.**
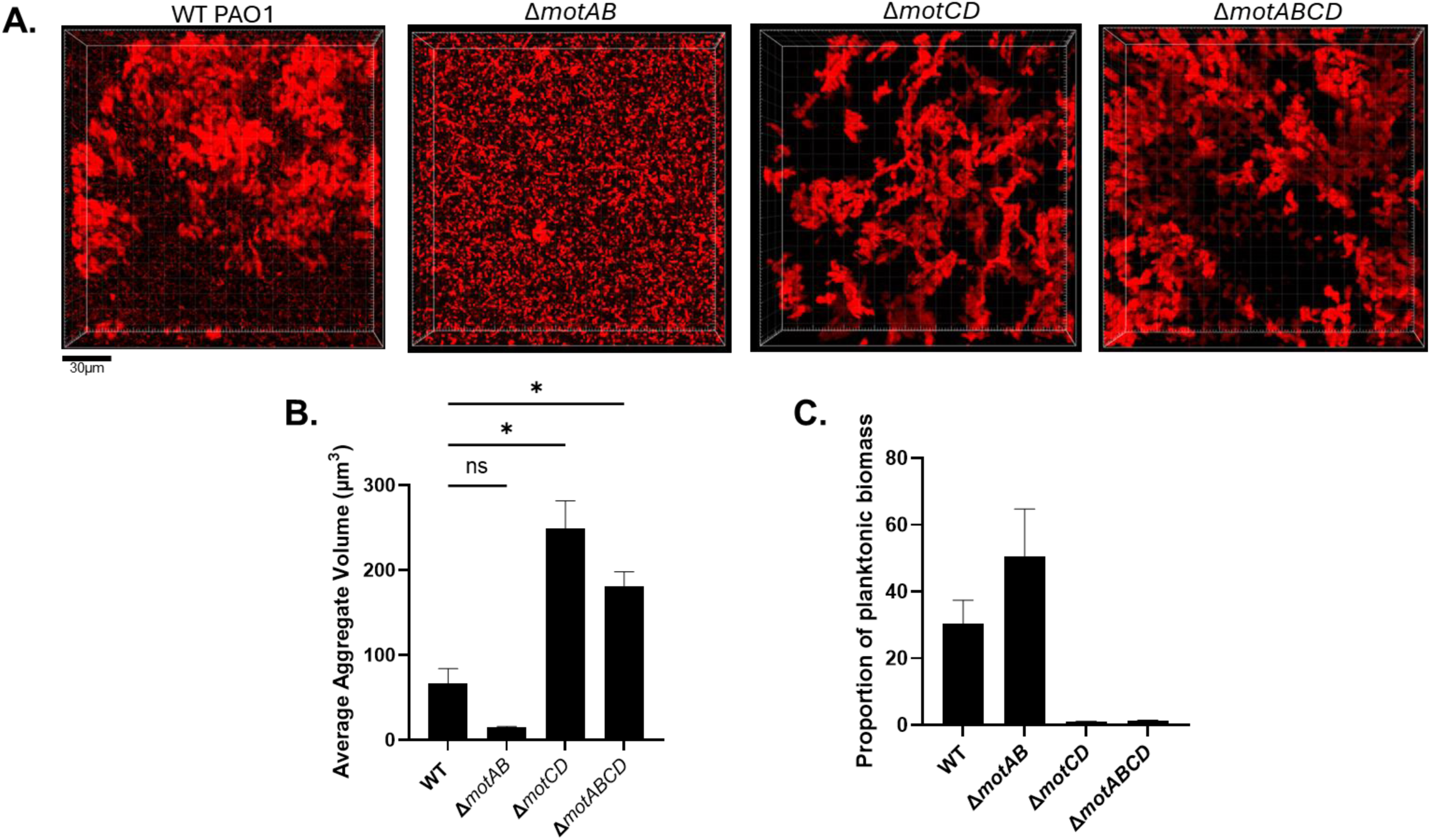
Mutants in flagellar rotation significantly impacts aggregation. Bacteria were grown in SCFM2 with 2% mucin for 6 hours before imaging. **A)** Representative images of flagellar mutants. Scale bar is 30μm. **B)** Quantification of aggregates **C)** Proportion of planktonic biomass of the indicated strains. *\*P<0.05* as determined by one-way ANOVA with Dunnett’s post-hoc test. Data shown are mean +/- SEM and are representative of 3 independent experiments. NS = not significant

Interestingly, we found an inverse correlation between antibiotic tolerance and surface biofilm formation where Δ*motAB* exhibited more biofilm formation than WT and Δ*motCD*, Δ*motABCD*, and Δ*fliC* all exhibited significantly less surface attached biofilm formation (**Fig. S4A**). These data support our initial assessment that aggregation is tightly linked to antibiotic tolerance and that the stator complexes have differential roles in aggregate formation in the diseased mucus environment.

### Motile subpopulations correlate with aggregative phenotypes

Several studies have investigated the differences between MotAB and MotCD stators in powering flagellar motility. While their functions are mostly redundant, they do have some unique properties. MotCD was found to be more important for swarming motility (33). Additionally, it was found that while MotAB provides more total rotational torque, MotCD provides better rotational stability (34). Therefore, we investigated the impact of the stator mutants on motility in our mucin rich system. As a first step, we assessed swimming motility using a traditional soft-agar based motility assay. We observed that both Δ*motAB* and Δ*motCD* exhibited a decrease in motility zones compared to WT mPAO1 (**Fig. S4B-C**). While this agreed with previous studies it did not explain the decreased tolerance and aggregation result for Δ*motAB* (**Figs. 2 and 3**). To better discern differences in the stator mutants, we used 2-D single cell motility tracking in our diseased mucus model. We grew bacteria in SCFM2 containing 2% mucin (w/v) for 1 hour prior to imaging. This timepoint was chosen to allow sufficient acclimation to the media, but also to retain a reasonable bacterial density for high resolution imaging of single cell motility. We observed multiple differences in motility behavior. Most notably, we found a significant difference in the proportion of motile bacteria between WT and the stator mutants. For WT mPAO1, we observed that ∼16% of the population was motile at a given time (**Fig. 4A-B**). The *motAB* mutant exhibited a significant increase in the proportion of motile bacteria. Conversely, less than 10% of *motCD* mutant cells were motile. Despite the significant difference in the motile subpopulation, there were only modest differences in the distance traveled (track length) of the motile bacteria between each strain (**Fig. 4A and C**). These data suggest that the proportion of motile bacteria within a population may directly correlate with aggregative capabilities.

**Figure 4.**
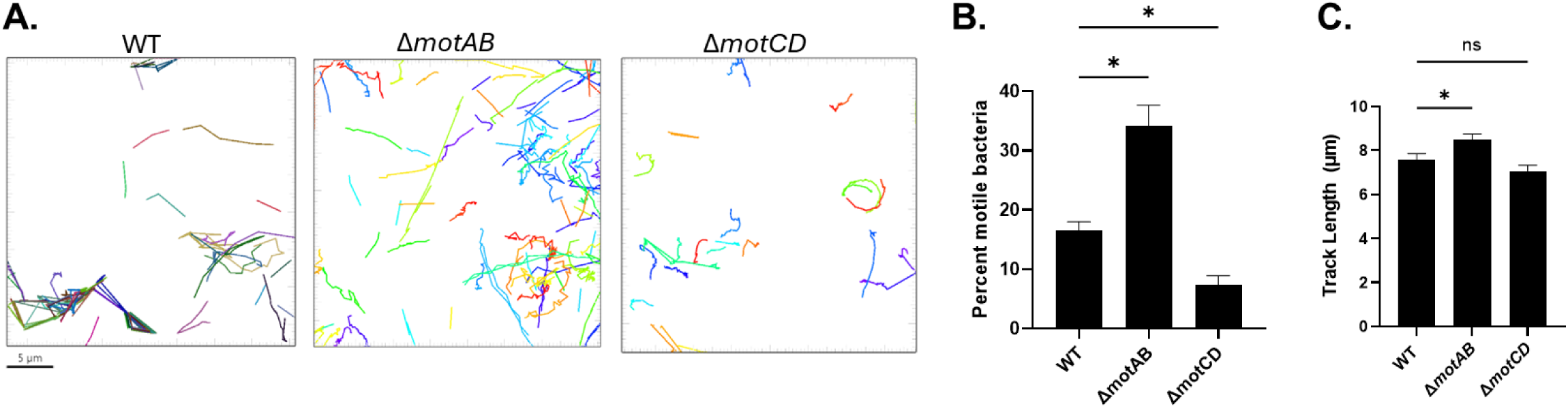
The MotAB and MotCD stators differentially contribute to motility in mucus. Exponential phase bacteria were inoculated into SCFM2 with 2% mucin for 1 hour. **A)** Representative traces of the motile bacteria from each strain, representing one field of view. Scale bar is 5μm. **B)** Quantification of the percent of motile bacteria within the tracked population. **C)** Average track length for the motile proportion. *\*P<0.05* as determined by one-way ANOVA with Dunnett’s post-hoc test. NS = not significant. Data shown are mean +/- SEM (**B and C**).

To better understand the direct impact of mucin on motility, we also conducted single cell motility tracking with bacteria grown in SCFM2 lacking mucin. In the absence of mucin, we observed a significantly higher proportion of motile bacteria (**Fig. S5**), indicating mucin polymers negatively impact motility.

### FliC regulation is important for tolerance and aggregation

Given our result that Δ*motAB* exhibited a significantly higher proportion of motile cells, we suspected that regulation of flagellar motility may also be important for aggregation. Downregulation of flagella is important for biofilm formation (35–37) and is likely also involved in aggregate formation. We engineered a strain that is incapable of shutting off flagellin expression and assessed whether the inability to control flagellin regulation impacted tolerance and aggregation. For constitutive *fliC* expression, we engineered the *fliC* gene such that expression was under the control of the synthetic IPTG inducible TAC promoter. We confirmed that both surface flagellin expression and swimming motility in soft agar is restored to levels comparable to WT (**Fig. S6**). When compared to WT in SCFM2 with 2% mucin, the constitutive *fliC* strain exhibited a marked reduction in tolerance to tobramycin (**Fig. 5A**) and decreased aggregation (**Fig. 5B-D**). We then reasoned that, similar to Δ*motAB*, a constitutive *fliC* strain may have a much higher proportion of motile cells within the population, which would hinder aggregation and therefore decrease tolerance. Indeed, using single cell tracking, we observed that the constitutive *fliC* strain exhibited a significantly higher proportion of motile cells compared to WT, as well as an increase in track length (**Fig. 5E-G**). These data suggest that regulation of flagella plays a key role in aggregate formation and antibiotic tolerance.

**Figure 5.**
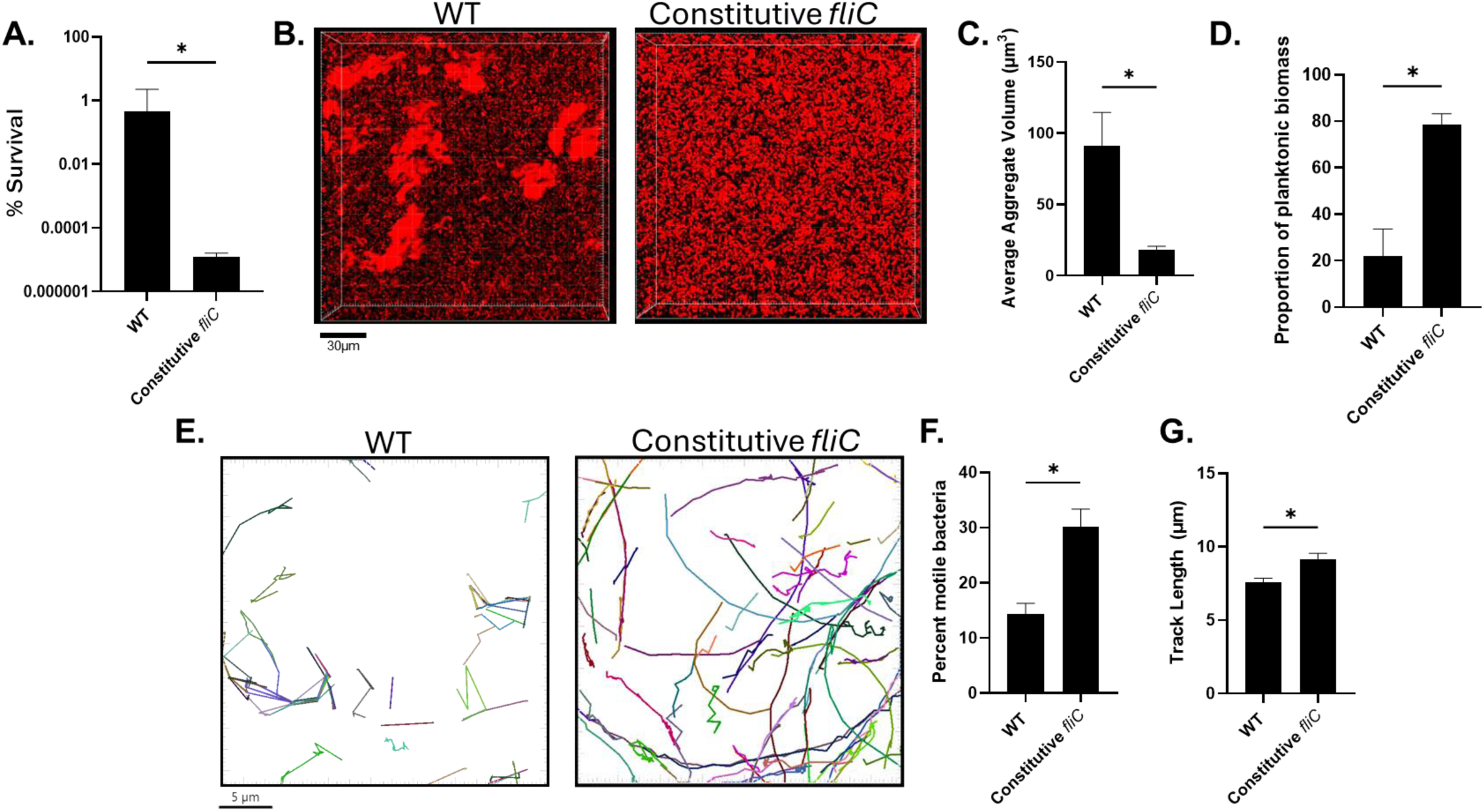
Constitutive *fliC* antagonizes tolerance and aggregation. **A)** Survival of WT or constitutive *fliC* expressing strain to tobramycin. **B)** representative images of aggregates after growth in 2% SCFM2 for 6 hours. Scale bar is 30μm. **C)** Quantification of aggregates from **B**. **D)** Proportion of planktonic biomass. **E)** Representative tracking traces and motility fraction proportions of WT or *fliC* constitutive strain as determined by single cell motility tracking. Scale bar is 5μm. **F)** Quantification of the percent of motile bacteria within the tracked population. **G)** Average track length for the motile proportion. *\*P<0.05* as determined by students *t*-test. Data shown are mean +/- SEM and are representative of 3 independent experiments (**A-C**), and a minimum of 800 cells tracked **(E-G).**

### Neutrophil elastase drives aggregation and antibiotic tolerance

Another hallmark of the MADs airway environment is the dominant neutrophil response. As a result, copious amounts of neutrophil effectors, particularly neutrophil elastase (NE), flood the airway. It has previously been shown that NE is capable of degrading flagellin of *P. aeruginosa* (38,39). Consequently, we evaluated if NE exposure would result in a similar phenotype as the flagellar mutants. We added disease relevant concentrations of NE (40) to SCFM2 containing 2% mucin and assessed tolerance to tobramycin and aggregation. We observed that exposure to NE led to an increase in tobramycin tolerance and an increase in aggregation (**Fig. 6A-C**). While the increase in aggregate size was not statistically significant, there was however a significant shift towards a reduced proportion of planktonic biomass in NE treated samples (**Fig. 6D**). Exposure to NE also led to a significant decrease in the proportion of motile bacteria and a decrease in track length (**Fig. 6E-G**). These data suggest that NE exposure drives antibiotic tolerance by impairing flagellar motility, thereby increasing aggregation. These observations suggest that while aggregate size likely plays a part in antibiotic tolerance, the proportion of bacteria in aggregates of any size (reduction of planktonic bacteria) has an important role in dictating antibiotic tolerance. Lastly, these data suggest that the host immune response may protect bacteria against antibiotic attack.

**Figure 6.**
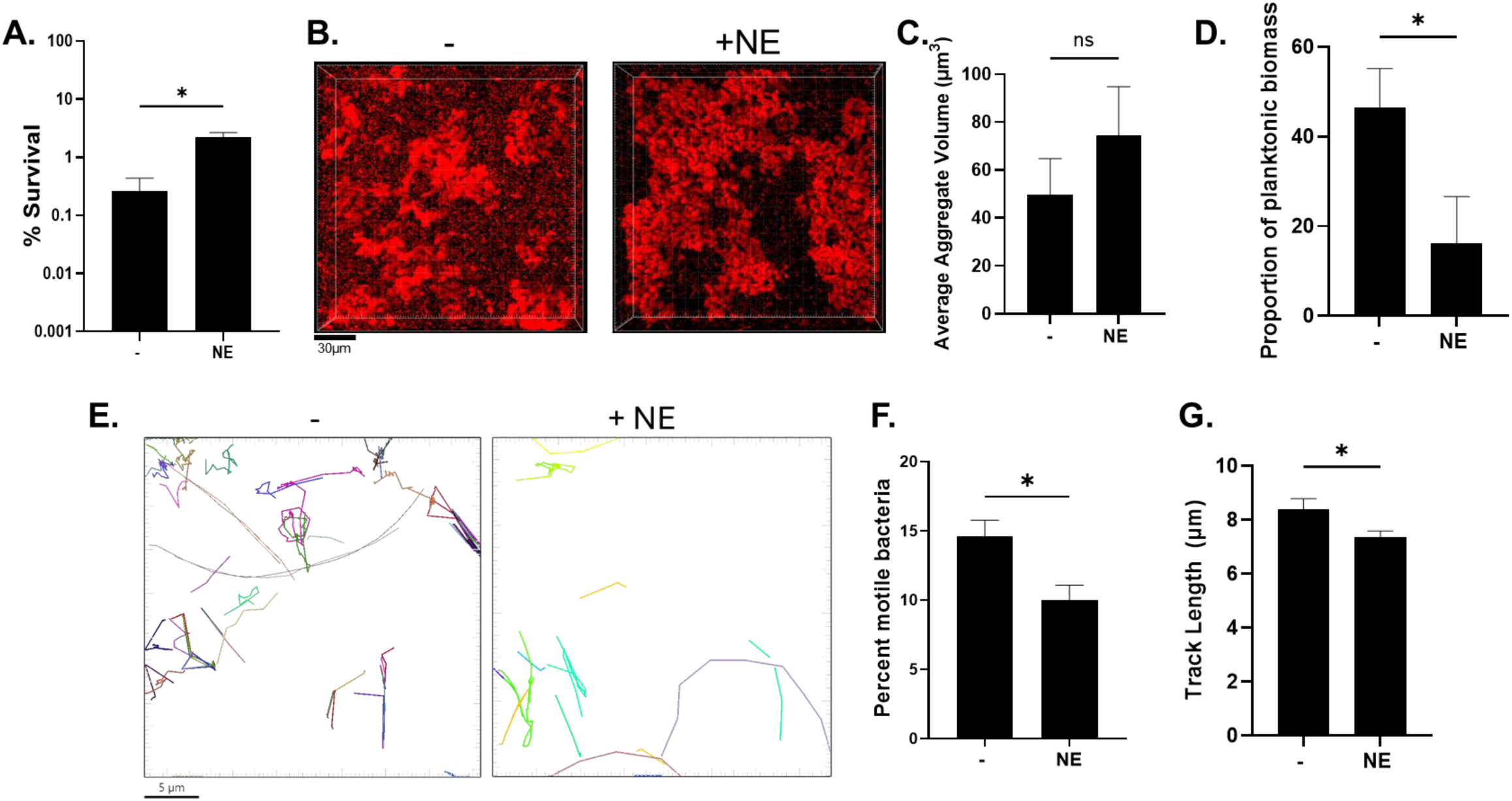
Neutrophil elastase drives aggregation and antibiotic tolerance. **A)** Bacteria were grown in normal SCFM2 media with 2% mucin (-) or with neutrophil elastase (+NE) (150μg/mL) for 8 hours then treated with tobramycin (300μg/mL) for 24 hours. **B)** representative images of aggregates after growth in 2% SCFM2 for 6 hours. Scale bar is 30μm. **C)** Imaris quantification of aggregates in (**B**). **D)** Proportion of planktonic biomass from analysis of (**B**). **E)** Representative tracking traces and motility fraction proportions of WT or *fliC* constitutive strain as determined by single cell motility tracking. Scale bar is 5μm. **F)** Quantification of the percent of motile bacteria within the tracked population. **G)** Average track length for the motile proportion. *\*P<0.05* as determined by students *t*-test. Data shown are mean +/- SEM and are representative of 3 independent experiments (**A-C**), and a minimum of 800 cells tracked **(E-G).**

## DISCUSSION

MADs are characterized by the accumulation of dehydrated mucus within the airways, which provides a niche for bacterial colonization and chronic infection. Chronic bacterial infections in MADs, where *P. aeruginosa* is the dominant pathogen, is the primary contributor to exacerbation, decreased quality of life, and early mortality. The failure of antibiotics to eradicate *P. aeruginosa* infections contributes significantly to mortality and morbidity in MADs (41–43). Understanding the mechanisms that contribute to antibiotic tolerance is critical to re-envisioning more effective therapies.

Our data show a strong correlation between the ability to form aggregates and antibiotic tolerance. Increases in mucin concentration led to increased antibiotic tolerance that was likely driven by the observed increase in aggregate size and decrease in planktonic biomass. Additionally, we showed that mucin polymers drive antibiotic tolerance by impeding flagellar motility and promoting the formation of aggregates. We speculate that when *P. aeruginosa* is trapped within the mucin polymer mesh, this likely facilitates cell-cell interactions thereby promoting aggregation. Alterations in flagellar motility that ultimately diminished motility (Δ*fliC*, Δ*flgE*, Δ*motCD*, Δ*motABCD*) in SCFM2 all exhibited both increased tolerance to antibiotics and increased aggregation. Notably, there was a marked decrease in the planktonic biomass when flagellar motility was diminished. It has been noted by many studies that planktonic bacteria are usually more susceptible to antibiotic treatments (44–47). The shift from the population from a planktonic to aggregated state is likely the primary driver of tolerance that we observed in this study.

The aggregates we observed were not adhered to a surface, but rather suspended in the media, and are more reminiscent of the aggregates observed in sputum, which are also not adhered to a traditionally defined surface. The *motAB* mutant, which exhibited lower tobramycin tolerance, exhibited increased biofilm, or surface attachment. This contradicts conventional logic in that better biofilm formation would confer tolerance. However, there are many stages of biofilm development and structure that likely impart various qualities that contribute to antibiotic tolerance. While the aggregates we observe are not adhered to a surface, they are likely to share many qualities of traditional surface attached biofilms. For instance, despite observations that *P. aeruginosa* resides in aggregate-biofilms that aren’t adhered to a traditional surface, it has been shown that, similar to biofilms, they produce exopolysaccharides (4). Additionally, despite flagellar mutants exhibiting lower biofilm, one study has shown that certain clinical isolates with defective flagellar motility exhibited increased production of Pel and Psl exopolysaccharides in a surface dependent manner (48).

Previous work has suggested that aggregate formation of *P. aeruginosa* in mucus is driven by a process termed “aggregation by depletion”, where entropy drives the formation of bacterial clusters in the presence of charged airway mucin and eDNA polymers (22,23). Here, we showed that loss of flagellar motility also drives aggregation. Though the mechanism of the observed increase in aggregation is still unknown, our results suggest that bacterial motility phenotypes, in combination with the environment, shape aggregative phenotypes and that aggregation likely occurs through active and passive processes. It is possible that some attachment structures, such as type IV pili, or other fimbria may be involved. Type IV pili have been described as a mediator of bacterial-mediated autoaggregation (49,50). However, it is difficult to discern fine differences between entropically driven aggregation and bacterial-mediated autoaggregation using our current system. It is also possible that loss of flagella, which renders bacteria non-motile, simply provides the opportunity of aggregation since the bacteria are unable to move away. Our results with the constitutive *fliC* strain, which was highly sensitive to tobramycin and did not form large aggregates, suggest that control of *fliC* expression is important for aggregate formation. It is possible that turning off flagellar motility is a response to becoming trapped in a polymer mesh, but the inability to control *fliC* expression allows bacteria to more readily escape mucin polymer entrapment, which would explain why the constitutive *fliC* strain is significantly more motile within SCFM2. Regardless, these results support the notion that there are ordered events that lead to the formation of aggregates.

While MotAB and MotCD have been described to have redundant functions, our results demonstrate that MotAB possesses a distinct role in aggregate formation. The MotCD complex has been shown to facilitate motility in higher viscosity environments (51). Therefore, when MotCD was absent (Δ*motCD*), in a viscous solution such as SCFM2 with 2% mucin, this likely resulted in the reduced motility, which then led to the increased aggregation and subsequently increased antibiotic tolerance. It has been shown that cells with only MotCD (i.e Δ*motAB*) had 10x more active motors than WT or cells with only MotAB (33). This could explain the phenomenon that Δ*motAB* had a higher proportion of motile cells which are less likely to become trapped by mucins.

One distinct function that has been described for MotAB is surface sensing. FimW is a c-di-GMP binding protein that localizes at cellular poles when contact with a surface occurs. This was shown to be dependent on MotAB (52). In a *motAB* mutant, there was a reduction in FimW localization, suggesting that MotAB is involved in surface sensing (52). It is possible that in our system, “surface sensing” can include non-traditional surfaces, like mucin polymers. We posit that sensing these non-traditional surfaces may be important for aggregation and that a *motAB* mutant, with defective surface sensing, is less efficient at forming aggregates.

Both phenotypic and genotypic heterogeneity is a hallmark of *P. aeruginosa* chronic infection in MADs (53,54). Heterogeneity is particularly relevant to antibiotic treatment failure, where a portion of a population possesses increased tolerance to antibiotics. Control of flagellar motility is a complex network of various regulators in which a diverse array of stimuli impacts motility (55). While complete ablation of flagellar biosynthesis is a common adaptation, other adaptations that alter flagellar motility without impacting flagellin expression, such as those affecting the motor/stator complex, the flagellar switch, or chemotaxis systems also arise (35,56,57). Our results help explain why defects in swimming motility are selected for over time during chronic infection as these mutations confer tolerance to multiple antibiotic classes. However, loss of flagellar motility may be detrimental in other aspects of infection. For instance, chemotaxis, either towards nutrients, or away from danger, relies on swimming motility. Without flagella, chemotaxis is limited and begs the question of how important chemotaxis is in chronic infection. It is possible that other forms of motility such as type IV pilus mediated twitching motility are used, though mutations in type IV pilus machinery and subsequently defects in twitching motility are common as well (58).

The inappropriate neutrophil response is a hallmark of MADs. Since *P. aeruginosa* primarily resides as aggregates within the airway, neutrophils often resort to releasing neutrophil extracellular traps (NETs) in order to deal with infection. One of the most abundant neutrophil effectors in NETs is neutrophil elastase (NE) (59–61), which also causes substantial damage to the host lung tissue. Leveraging previous literature and based on our results with flagellar mutants, we had predicted that exposure to NE would increase aggregation and tolerance through its effect on flagella. Indeed, we showed that NE decreased flagellar motility and subsequently increased aggregation and tolerance to tobramycin. These results suggest that a novel strategy to limit tolerance could be to use an already FDA approved NE inhibitor such as Sivelestat. While NE inhibitors are sparingly used, their main indication is to target inflammation. However, using NE as a new indication for treating antibiotic tolerant infections could be presented as a novel therapeutic in conjunction with tobramycin therapy.

Collectively, our results show a strong correlation between aggregate formation and antibiotic tolerance of *P. aeruginosa* in an *in vitro* mimic of the MADS lung environment. Adaptation to host environmental factors shifts *P. aeruginosa* into a more aggregative state, thereby conferring tolerance to multiple classes of antibiotics. Flagellar motility plays a key role in the formation of aggregates within the diseased mucus environment and alterations of motility can skew population aggregation phenotypes. Host immune derived factors such as NE negatively impact motility and help drive antibiotic tolerance. The results of this study shed light on how such a common adaptation is advantageous in the presence of antibiotics and also deepens our understanding of why mutants in flagellar motility are selected for during chronic infection.

## MATERIALS AND METHODS

### Bacterial Strains and Culture Conditions

All bacterial strains and plasmids used in this study are listed in **Table S1**. Bacteria was swabbed from frozen stocks onto Lysogeny Broth (LB) (Miller) agar and incubated overnight at 37°C. Overnight liquid cultures were inoculated from single colonies and were shaken overnight in LB broth at 250 RPM at 37°C. SCFM2 was prepared as described (21,24), or purchased from Synthbiome. Where indicated, antibiotics or neutrophil elastase were added. Neutrophil elastase (Innovative Research) was added at 150μg/ml.

### Generation of mutants, complementation, and constitutive expression plasmids

Primers used in this study are listed in **Table S2**. Deletion of genes was achieved through SacB assisted allelic exchange, using the Gateway Cloning (GW) platform (Invitrogen) (62). Fluorescent bacteria were generated via triparental mating of *P. aeruginosa* with *E. coli* containing the pUC18T-mini-Tn7T-Gm-Pc-DsRed- Express2 (26) plasmid and *E. coli* containing the pTNS2 helper plasmid (63). Vector backbone was removed through flp recombinase (64). Complementation of *fliC* was achieved through amplifying the *fliC* gene and its promoter region (500bp upstream of start codon), then using GW to introduce *fliC* and its promoter into a GW adapted pMini-CTX vector for chromosomal complementation at the neutral ΦCTX site. The vector backbone was removed through flp recombinase (64). Complementation was confirmed by PCR and swimming motility assays. Constitutive expression of *fliC* was achieved by amplifying the *fliC* gene and introducing it via GW cloning into the pMMB67 vector, which contains the TAC promoter that is inducible by isopropyl β-D-1- thiogalactopyranoside (IPTG). Expression of *fliC* was achieved through addition of 100μM IPTG to culture conditions. IPTG was only added during culture in SCFM2, and not during overnight culture.

### Antibiotic susceptibility assays

Antibiotic susceptibility assays were performed as previously described (21). Briefly, overnight cultures of *P. aeruginosa* were subcultured into fresh LB at a 1:50 dilution and cultured to exponential phase until an OD_600_ of 0.25 was achieved. Exponential phase bacteria were then inoculated 1:100 into SCFM2 for an inoculum of 1×10^6^ CFU/mL. Bacteria were then incubated statically at 37°C for 8 hours. At 8 hours, duplicate wells were collected and serial diluted and plated for CFU at time of treatment (“At Treats”). Another set of wells were then treated with various antibiotics: tobramycin at 300μg/ml, ceftazidime/avibactam at 1000/40 μg/ml, and meropenem at 2000μg/mL. All antibiotics were purchased through Sigma Aldrich. Treatment went for 24 (standard assay) or up to 72 hours (persistence assays) before bacteria were collected, washed twice in M63 salts, and plated for enumeration. Percent survival was calculated using the following:

### Swimming motility assays

Bacteria were grown overnight in LB and subcultured for 1 hour at a 1:50 dilution. 2μl of subculture was inoculated into LB +0.3% agar. Plates were incubated at room temperature for 30 hours before the zone of motility was measured. Imaging of swimming plates was achieved using the iBright FL1500 imager (Applied Biosystems) using the 490-520 (TRANS) filter in the visible channel.

### Fluorescence microscopy

Bacteria were prepared the same as for antibiotic survival assays described above. 1×10^6^ CFU/mL was inoculated into 300 μL SCFM2 in 8-well #1.5 coverglass bottom chamber slides (Cell-Vis, Cat# C8-1.5P). After 6 hours of static incubation at 37C, the center of the wells were imaged using a Leica Stellaris5 laser scanning confocal microscope with an environmental box (Okolab) set to 37C for live cell imaging. Fluorescence was observed with a white light laser at a laser line of 554nm at 5% power and detection range of 569-650nm and gain of 60. Using a 63x, HC PL APO CS2 oil immersion objective with a numerical aperture of 1.4 and a pinhole diameter of 1 Airy Unit (AU), we obtained 180×180×30 μm 3D Z-stacks at a resolution of 1024×1024 and scan speed of 600hz, beginning at least 10μm above the bottom of the glass. Images were captured using the LAS X software version 4.5 (Leica microsystems). Raw Leica images files (LIF) were then exported for analysis.

### Motility Tracking

Motility tracking was achieved with the same methods as described above for fluorescent microscopy with some modifications. Bacteria were inoculated into 100μL SCFM2 at 5×10^6^ CFU/mL and grown statically for 1 hour prior to imaging. 2D videos of 30.75×30.75μm were captured at 256×256 resolution with a zoom of 6 and bidirectional scanning at a speed of 1600hz (∼11.76 frames per second). Pinhole diameter was set to 6.28 AU. Videos were approximately 1 minute long. A minimum of 10 videos were captured per condition/strain and a minimum of 800 cells were tracked. Video files were then exported for analysis.

### Image Analysis

Analysis of aggregates and motility tracking were performed using Imaris 10.0 (Oxford Instruments). LIF files were imported into Imaris. Images were visualized in Imaris as a 3D max projection. For analysis of aggregates, we created a custom surfaces creation parameter algorithm. Background subtraction threshold minimum value was set to 15. Surfaces with volumes less than 0.335μm^3^ or greater than 5000μm^3^ were filtered out as artifacts. Surfaces with volumes greater than 5μm^3^ were considered aggregates. Examples of aggregates are highlighted in **Fig. S1A**. Planktonic biomass proportion was determined as the ratio of the sum of surface volumes between 0.335-5μm^3^ to the sum of all surfaces calculated.

For motility tracking, we used the spots function to track bacterial motility. Minimum quality threshold was set to 30 and maximum gap size was set to zero. For motile bacteria, we set a track distance minimum of 2μm and track duration between 0.050-5.00 seconds. Traces of motile bacteria were highlighted in Imaris and exported.

### Immunoblot of FliC

Visualization of FliC was achieved through western blotting. Strains were inoculated into 2% mucin SCFM2 and grown for 8 hours. For isolation of surface flagella, bacteria were collected and centrifuged at 7,500x *g* for 3 minutes to pellet the bacteria without shearing flagella. The supernatant was removed, and pellets resuspended in 150 μl M63 salts. Using a 1mL syringe and 25-gauge needle, the suspensions were passed through the needle 20 times. The cells were pelleted at 17,000 x *g* for 2 minutes, and the supernatant containing sheared flagella from the cell surface was collected. Sample protein content was normalized by quantifying protein content of the cell pellets using the bicinchoninic acid (BCA) assay (Thermo Fisher). Samples were adjusted according to cell pellet protein content and ran on a pre-cast 10% TGX acrylamide gel (Bio-Rad) with the Precision Plus Protein Dual Color Standards ladder (Bio-Rad) for 1.5 hours at 80 volts. Proteins were then transferred to a PVDF membrane using standard wet transfer at 90 volts for 1 hour at 4C. Following blocking using 5% milk for 1 hour, the membrane was probed with rabbit anti-FliC polyclonal antibody (65) (1:2000) overnight at 4C, followed by donkey anti-rabbit secondary antibody conjugated to IRDye680 fluorophore (1:20,000; LI-COR Biosciences, Cat# 926-68023) for 2 hours. Bands were visualized on an iBright FL1500 imager using the X4 (610-660nm excitation) and M4 (710-730nm emission) pre-configured filter set for IRDye 680.

### Biofilm assay

Biofilm assays were performed using the crystal violet staining method, as previously described (66) Briefly, overnight cultures of bacteria were subcultured at 1:50 dilution for 1 hour in LB to allow bacteria to enter exponential phase. 1×10^6^ CFU/mL were then inoculated into SCFM2 containing 2% mucus in tissue-culture treated 96-well plates. Bacteria were incubated statically at 37C for 24 hours. To measure biofilm, after incubation, plates were washed in water and dried for 2 hours in ambient air before adding 0.1% crystal violet to stain attached bacteria for 15 minutes at room temperature. After staining, plates were washed 3 times in water and allowed to dry overnight. 95% ethanol was then added to the wells and the plate was incubated for 15 minutes at room temperature. Well contents were transferred to a clean 96-well plate and the absorbance at 550nm was measured using a Tecan Infinite M Plex plate reader. Wells containing sterile media were used as blank and negative controls.

### Statistical Analysis

All experiments were performed in at least biological triplicate and across different days and media preparations. Statistical analysis was achieved via student’s two-tailed *t-*test, one way analysis of variance (ANOVA), or two-way ANOVA as indicated. Differences were considered significantly different with *P-*value <0.05. Statistical tests were carried out using GraphPad Prism version 10.2.

### Data Availability

Custom Imaris creation parameters used for the analysis of aggregates and motility tracking are publicly available in the Carolina Digital Repository at: https://doi.org/10.17615/tvvm-8f83.

## Acknowledgements

We thank Dr. Stephen Lory at Harvard Medical School for providing the FliC antibody. We thank Dr Michael Chua in the Michael Hooker Microscopy Facility at UNC Chapel Hill for assistance in confocal imaging. We thank Dr. Michelle Itano in the Neuroscience Microscopy Core at UNC Chapel Hill for the use of and their assistance with Imaris. Microscopy analysis was performed at the UNC Neuroscience Microscopy Core (RRID:SCR_019060), supported, in part, by funding from the NIH-NICHD Intellectual and Developmental Disabilities Research Center Support Grant P50 HD103573.

## Supplementary Material

### Supplementary Figures

**Figure S1.**
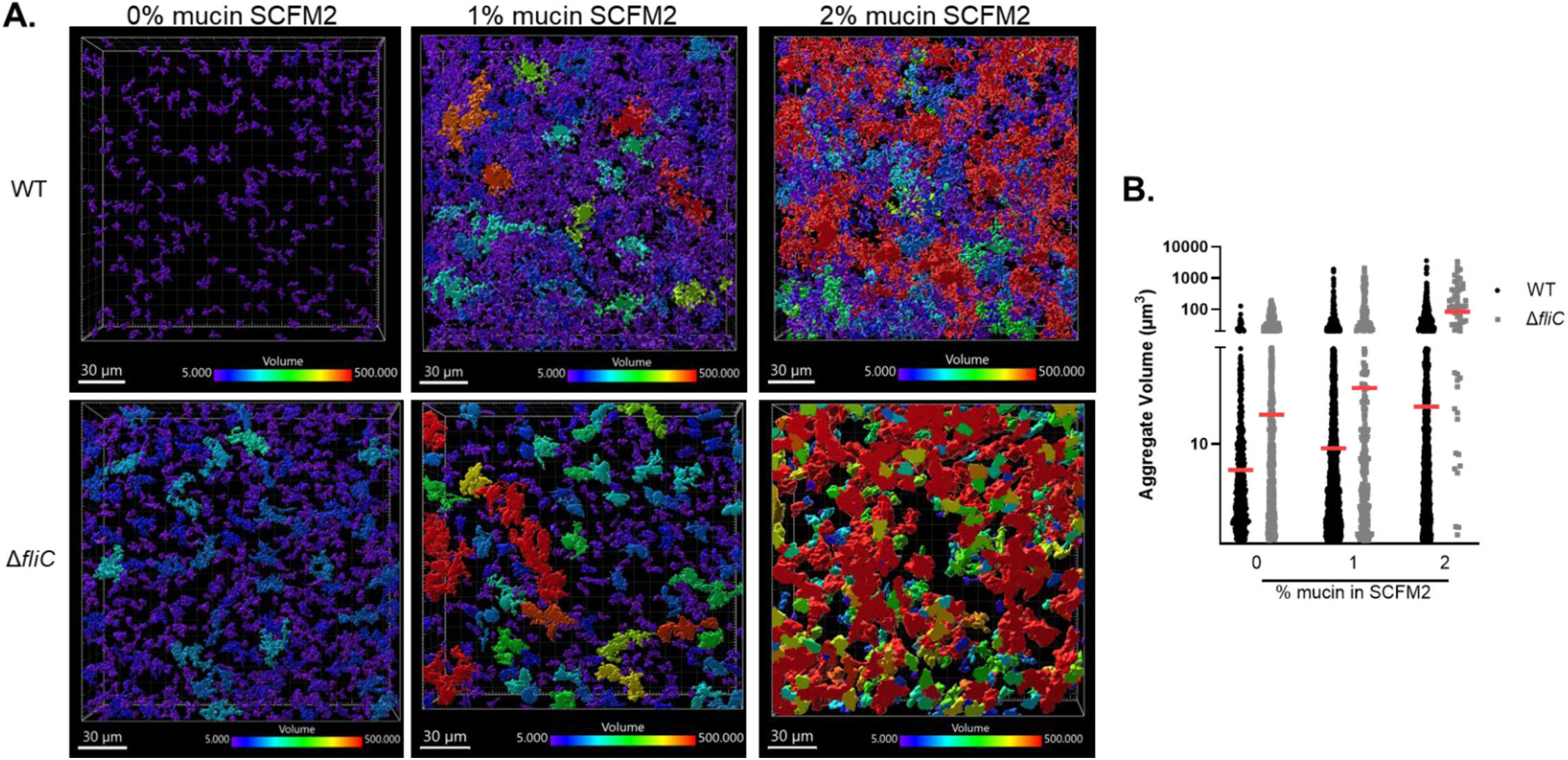
Visualization of Imaris calculated aggregates. **A)** Representative images from Figure 1 with individual aggregate surfaces highlighted using Imaris. Scale ranges from 5-500μm^3^. **B)** Aggregate distribution by size from a single image.

**Figure S2.**
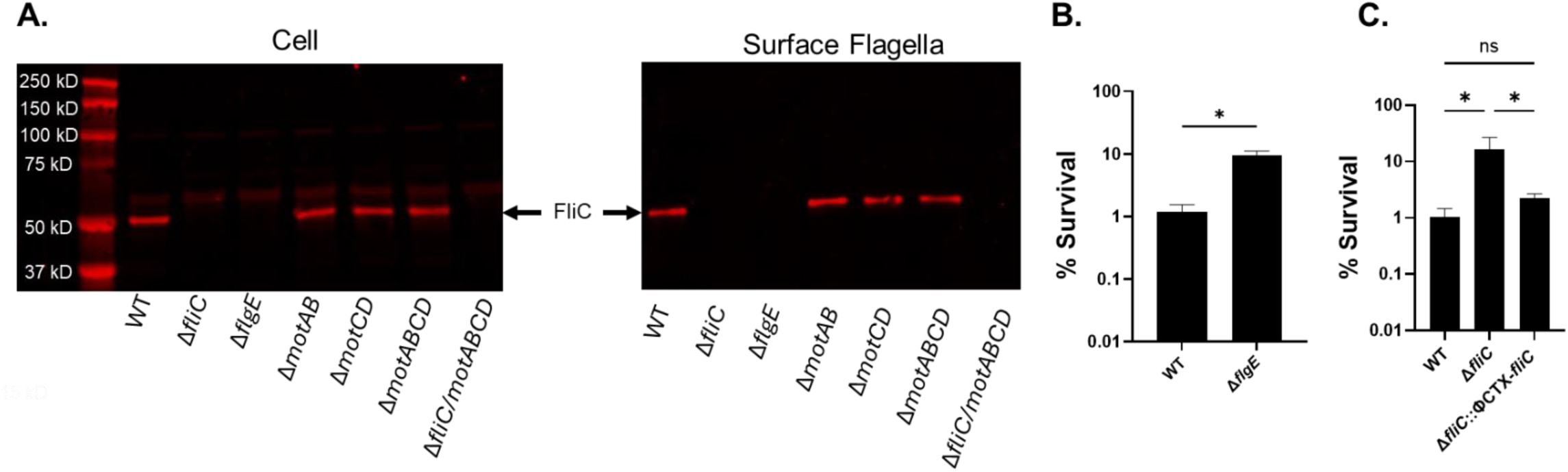
*fliC* complementation restores tolerance and motor protein mutants still produce surface flagella. **A)** Bacteria were grown in SCFM2 containing 2% SMM for 8 hours. Cell and surface fraction were probed for FliC via western blot **B)** WT PAO1 and Δ*flgE,* and **C)** WT PAO1, Δ*fliC*, or *fliC* native complement (Δ*fliC::*ΦCTX-*fliC*) were grown in SCFM2 with 2% mucin for 8 hours, then treated with tobramycin (300μg/mL) for 24 hours. Percent survival is plotted as mean +/- SEM. *\*P<0.05,* as determined by **B)** unpaired t-test or **C)** one way ANOVA with Dunnett’s post hoc test.

**Figure S3.**
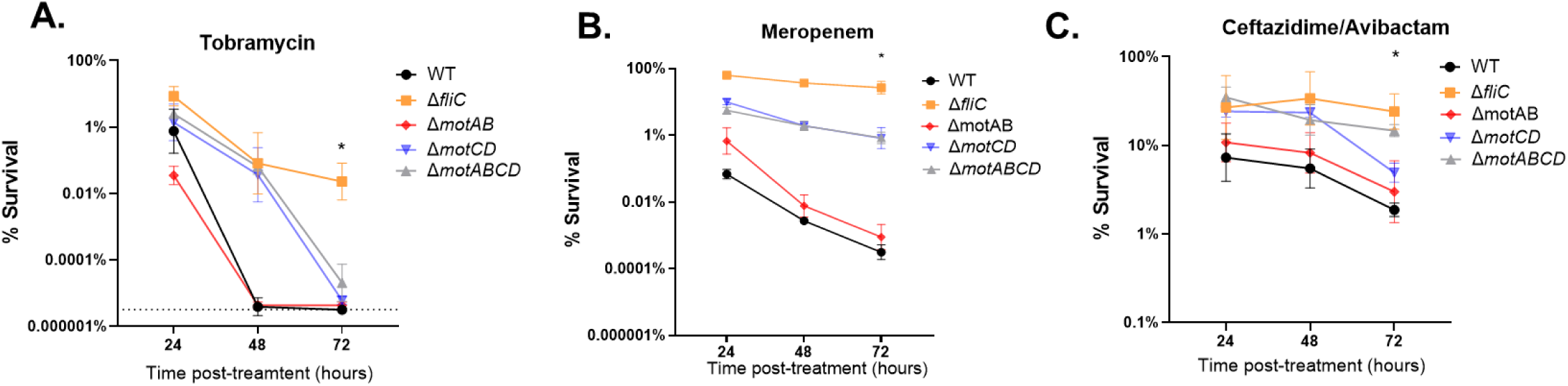
Loss of flagellar motility increases tolerance and persistence to multiple classes of antibiotics. Bacteria were grown for 8 hours in 2% mucin SCFM2 then treated with **A)** tobramycin (300μg/mL), **B)** meropenem (2000μg/mL), or **C)** ceftazidime/avibactam (1000/40 μg/mL) for up to 72 hours. Dashed line in (**A**) indicates limit of detection.

**Figure S4.**
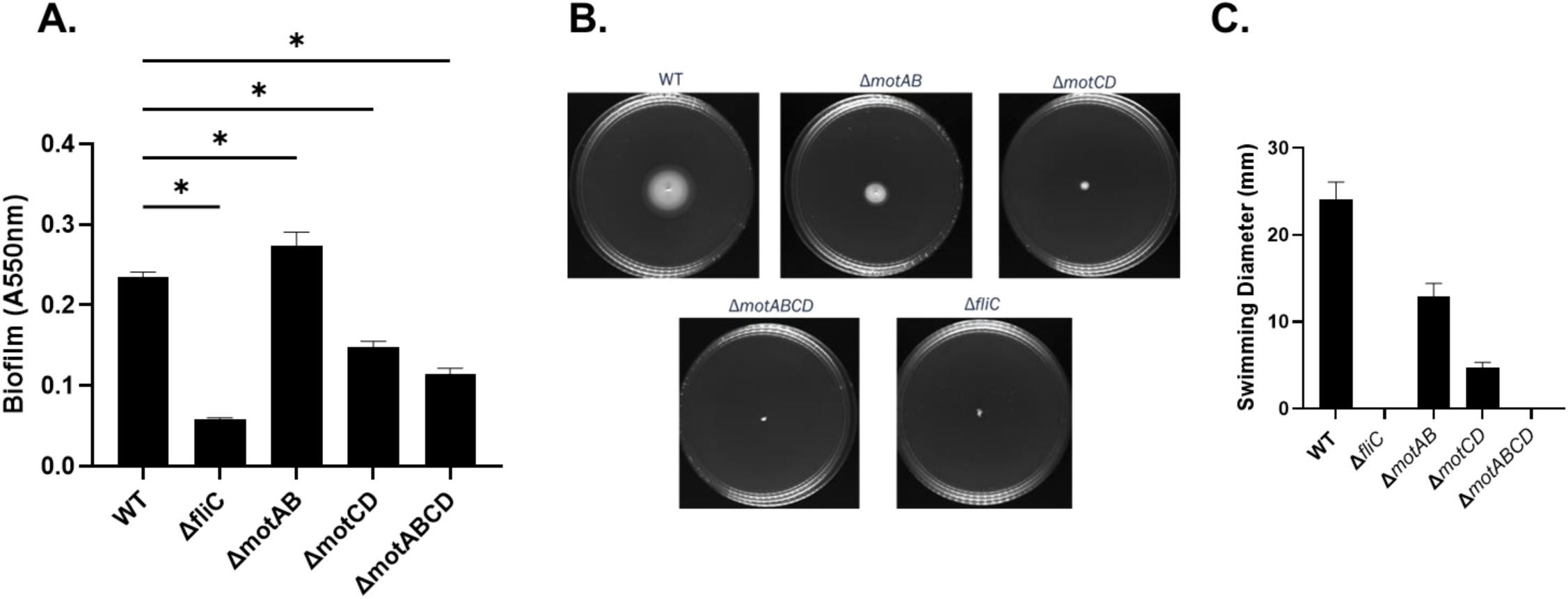
Biofilm formation and swimming motility of flagellar mutants. **A)** Bacteria were grown in 2% mucin SCFM2 for 24 hours before staining surface attached bacteria with 0.1% crystal violet. Absorbance was measured at 550nm. **B)** Bacteria were inoculated into LB with 0.3% agar and incubated at room temperature for ∼30 hours before zone of motility was measured. Representative images of motility zones shown. **C)** Quantification of swimming motility from (**B**).

**Figure S5.**
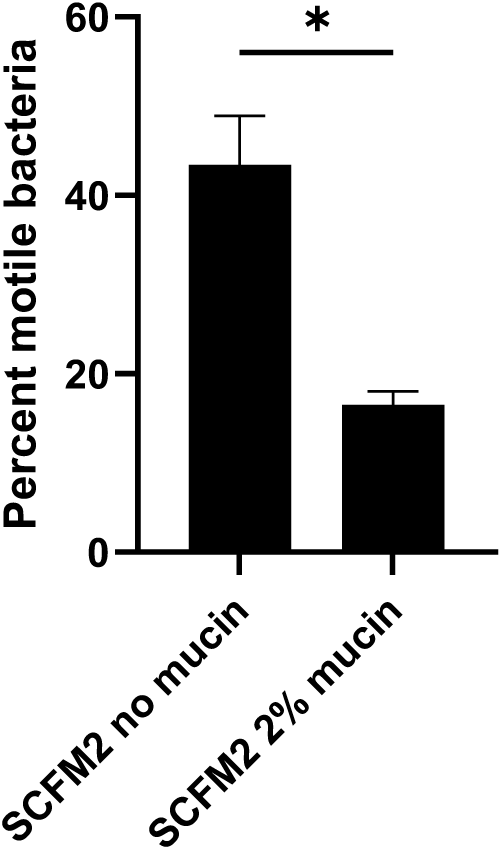
Mucin constrains motility. Single cell motility tracking was conducted either in SCFM2 with 2% mucin or SCFM2 lacking mucin. The percent of motile bacteria is presented as mean +/- SEM. \**P<0.05* as determined by student’s *t-*test

**Figure S6.**
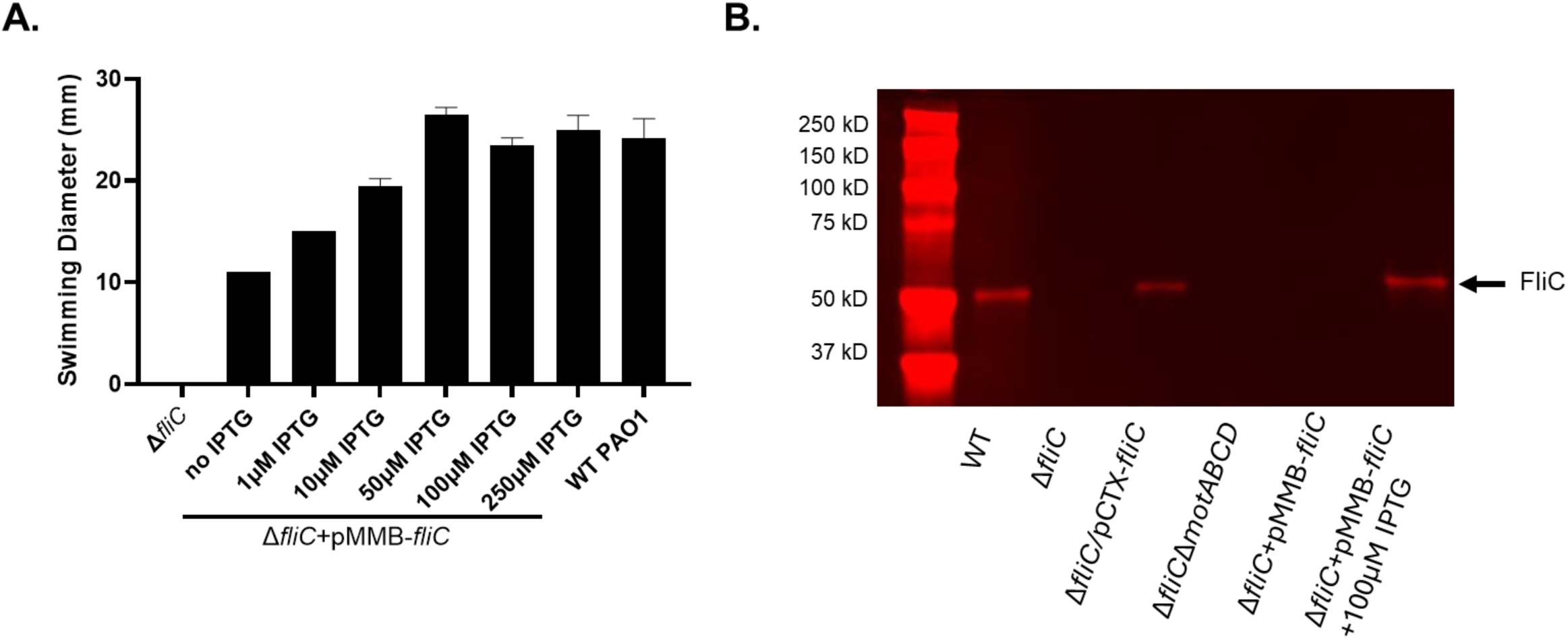
Inducible *fliC* expression restores flagellin levels and swimming motility. **A)** Swimming motility in soft agar for WT and constitutive *fliC* strain with varying concentrations of IPTG. 100μM IPTG was chosen for subsequent experiments. **B)** Western blot of surface flagella of indicated strains.

**Table S1:** Strains and plasmids used in this study.

| Strain/plasmid | Description | Source/Reference |
| --- | --- | --- |
| *Pseudomonas aeruginosa* |  |  |
| mPAO1 | WT mPAO1 | (67) |
| $\Delta fliC$ | In frame deletion of *fliC* | This study |
| $\Delta flgE$ | In frame deletion of *flgE* | This study |
| $\Delta fliC + \Phi CTX - fliC$ | Native *fliC* complement via insertion of *fliC* and promoter into neutral $\Phi CTX$ site | This study |
| $\Delta motAB$ | In frame deletion of *motAB* | This study |
| $\Delta motCD$ | In frame deletion of *motCD* | This study |
| $\Delta motABCD$ | In frame deletion of *motABCD* | This study |
| WT mPAO1 dsRed-Express2 | WT mPAO1 expressing DsRed-express2 from Tn7 site | This study |
| $\Delta fliC$ dsRed-Express2 | $\Delta fliC$ strain expressing DsRed-express2 from Tn7 site | This study |
| $\Delta motAB$ dsRed-Express2 | $\Delta motAB$ strain expressing DsRed-express2 from Tn7 site | This study |
| $\Delta motCD$ dsRed-Express2 | $\Delta motCD$ strain expressing DsRed-express2 from Tn7 site | This study |
| $\Delta motABCD$ dsRed-Express2 | $\Delta motABCD$ strain expressing DsRed-express2 from Tn7 site | This study |
| Constitutive *fliC* | $\Delta fliC$ strain carrying pMMB plasmid with *fliC* under TAC promoter | This study |
| Constitutive *fliC* dsRed-express2 | Constitutive *fliC* strain expressing dsRed-Express2 at Tn7 site | This study |
| **Plasmids** |  |  |
| pDONR201 | Gateway entry vector | Invitrogen |
| pEXG2 | Suicide plasmid for in frame deletions | (68) |
| pFLP2 | Vector containing flp recombinase for removing vector backbone of CTX and Tn7 plasmids | (64) |
| pTNS2 | Helper plasmid for transformation of pUC18-mini-Tn7 vectors | (63) |
| pMMB67 | IPTG inducible Expression vector | (69) |
| pMMB-*fliC* | pMMB67 with *fliC* ORF under TAC promoter | This study |
| pUC18T-mini-Tn7T-Gm-Pc-DsRed-Express2 | Tn7 vector for insertion of DsRed-express2 fluorophore | (70) |
| pMini-CTX | CTX vector for chromosomal insertion | (71) |

**Table S2:**
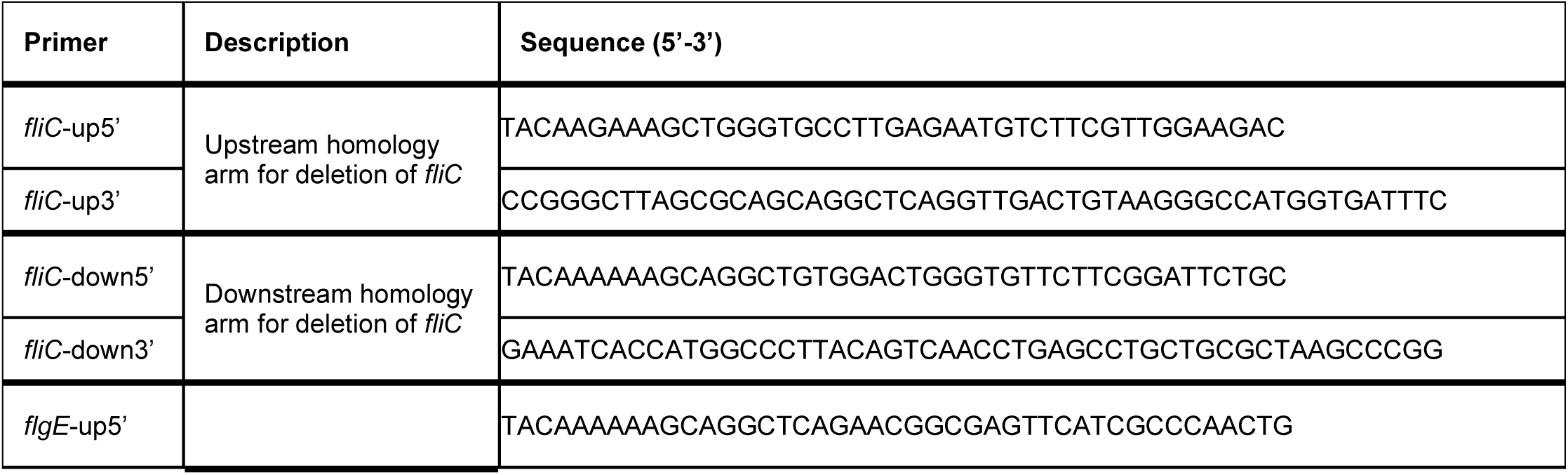
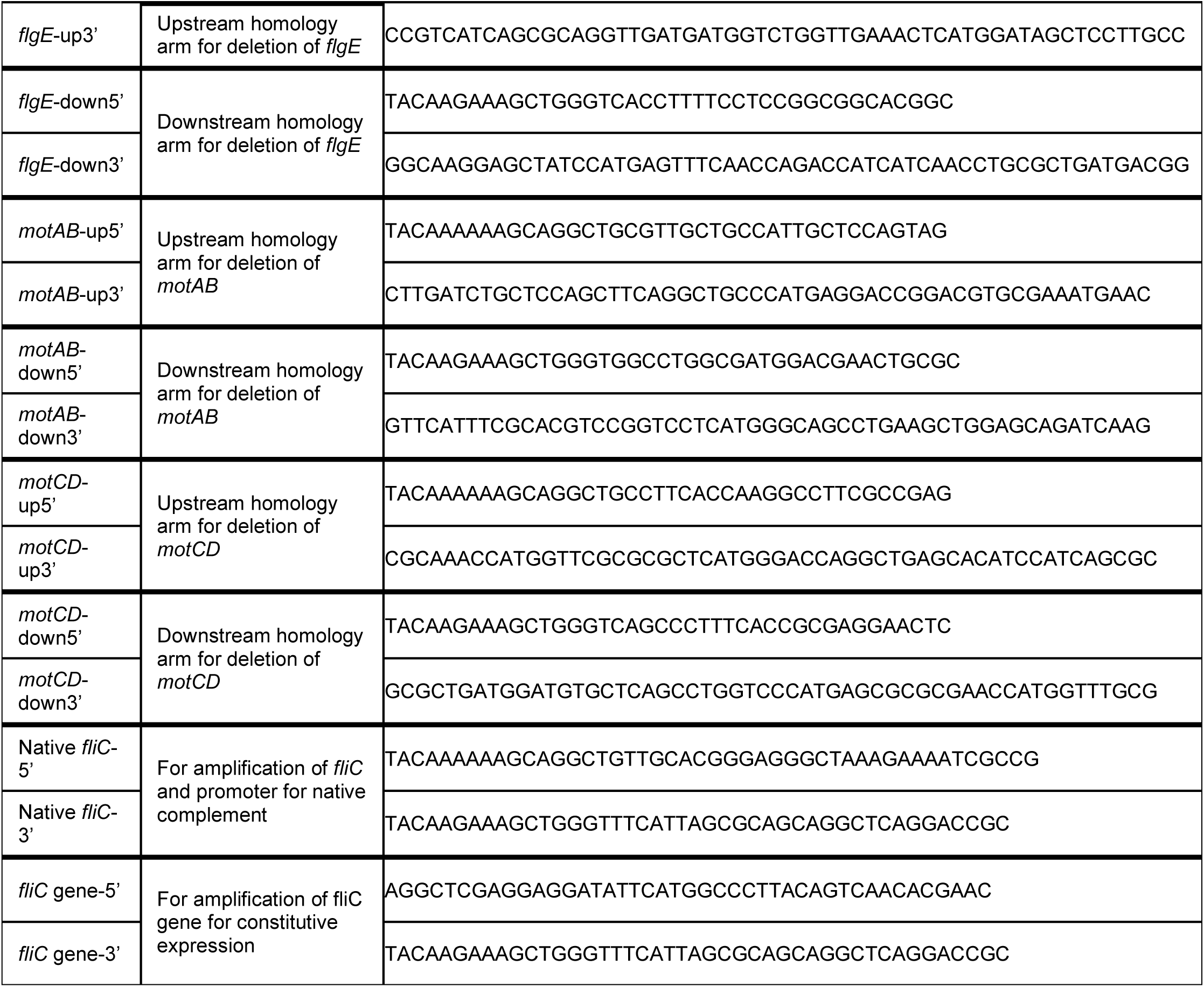
Relevant primers used in this study.

